# Addressing challenges of high spatial resolution, UHF field fMRI for group analysis of higher-order cognitive tasks; an inter-sensory task directing attention between visual and somatosensory domains

**DOI:** 10.1101/373977

**Authors:** Kevin M. Aquino, Rodika Sokoliuk, Daisie Pakenham, Rosa Sanchez Panchuelo, Simon Hanslmayr, Stephen D. Mayhew, Karen J. Mullinger, Susan T. Francis

## Abstract

Functional MRI at ultra-high field (UHF, ≥7T) provides significant increases in BOLD contrast-to-noise ratio (CNR) compared with conventional field strength (3T), and has been exploited for reduced field-of-view, high spatial resolution mapping of primary sensory areas. Applying these high spatial resolution methods to investigate whole brain functional responses to higher-order cognitive tasks leads to a number of challenges, in particular how to perform robust group-level statistical analyses.

This study addresses these challenges using an inter-sensory cognitive task which modulates top-down attention at graded levels between the visual and somatosensory domains. At the individual level, highly focal functional activation to the task and task difficulty (modulated by attention levels) were detectable due to the high CNR at UHF. However, to assess group level effects, both anatomical and functional variability must be considered during analysis. We demonstrate the importance of surface over volume normalization and the requirement of no spatial smoothing when assessing highly focal activity. Using novel group analysis on anatomically parcellated brain regions, we show that in higher cognitive areas (parietal and dorsal-lateral-prefrontal cortex) fMRI responses to graded attention levels were modulated quadratically, whilst in visual cortex and VIP, responses were modulated linearly. These group fMRI responses were not seen clearly using conventional second-level GLM analyses, illustrating the limitations of a conventional approach when investigating such focal responses in higher cognitive regions which are more anatomically variable. The approaches demonstrated here complement other advanced analysis methods such as multi-variate pattern analysis, allowing UHF to be fully exploited in cognitive neuroscience.

## Introduction

The development of ultra-high field (UHF, ≥ 7T) MRI scanners has provided new opportunities for functional MRI (fMRI). Increasing the field strength results in the intrinsic increase in image signal-to-noise ratio (SNR) (Vaughan, Garwood et al. 2001, Pohmann, Speck et al. 2016) and this, coupled with an increased blood oxygenation level-dependent (BOLD) signal change, results in increased BOLD contrast-to-noise ratio (CNR). This can be exploited to improve the spatial resolution of fMRI data or to enhance the sensitivity, enabling the detection of weaker responses.

To date, the majority of UHF fMRI studies have used reduced field-of-view (FOV) data acquisitions to study chosen primary sensory areas, such as the visual and sensorimotor cortices (Fracasso, Luijten et al. 2017, Reithler, Peters et al. 2017, Schluppeck, Sanchez-Panchuelo et al. 2017), thus overcoming a number of challenges associated with B_0_ and B_1_ inhomogeneities in larger FOV acquisitions (Polimeni, Renvall et al. 2017, Uludag and Blinder 2017). For example, the increase in BOLD CNR of UHF experiments has been used to provide detailed maps of individual subjects’ visual (Goncalves, Ban et al. 2015, Kemper, De Martino et al. 2017, Poltoratski, Ling et al. 2017, Rua, Costagli et al. 2017) and somatosensory functional responses (Sanchez Panchuelo, Schluppeck et al. 2015, Sanchez Panchuelo, Ackerley et al. 2016) and how these relate to individual brain anatomy (Sanchez-Panchuelo, Besle et al. 2012, Besle, Sanchez-Panchuelo et al. 2014, Sanchez-Panchuelo, Besle et al. 2014). These functional maps have been shown to spatially vary across subjects, highlighting inter-subject variability, whilst the intra-subject session reproducibility of these maps has been shown to be high (Sanchez-Panchuelo, Besle et al. 2012, Goncalves, Ban et al. 2015). Imaging at the sub-millimetre level has allowed mapping of cortical columns and “layers” of cortex (e.g. (Yacoub, Shmuel et al. 2007, Zimmermann, Goebel et al. 2011, Olman, Harel et al. 2012, De Martino, Moerel et al. 2015, Muckli, De Martino et al. 2015, Kok, Bains et al. 2016)), providing a novel method by which to distinguish bottom-up and top-down neural processes (Olman, Harel et al. 2012, Muckli, De Martino et al. 2015, Kok, Bains et al. 2016).

The challenges associated with increased image distortion in full FOV acquisitions at UHF have limited the study of whole brain cortical function at high spatial resolution. Further, large inter-individual anatomical differences can arise (Geyer, Weiss et al. 2011, Kanai and Rees 2011, Gu and Kanai 2014), which result in challenges in the data analysis of group functional responses, meaning such responses might be lost at the group level due to lack of spatial congruency. Only a limited number of publications have studied whole brain functional responses at UHF (e.g. (Boyacioglu, Schulz et al. 2014, Vu, Phillips et al. 2016, Goodman, Wang et al. 2017, Mestres-Misse, Trampel et al. 2017)), with the study of cognitive function being limited, as highlighted in a recent review article (De Martino, Yacoub et al. 2017). To date, to our knowledge, only two studies have used UHF fMRI to map responses to complex cognitive tasks in higher-order cortical regions over the whole brain (Vu, Phillips et al. 2016, Goodman, Wang et al. 2017). Goodman *et al* (Goodman, Wang et al. 2017) exploited 7T to investigate the neural basis of consumer buying motivations. This study maximised the increased BOLD CNR of UHF, but did not realise the full potential of 7T as a 6mm full-width-at-half-maximum (FWHM) smoothing kernel was applied to GE-EPI data acquired at 2 mm isotropic to reveal the group activity. Vu *et al* (Vu, Phillips et al. 2016) exploited the benefits of high 500 ms temporal resolution to show the improved sensitivity of capillary responses at 7T compared to 3T allowing the decoding of fine-grained temporal information for word classification.

UHF provides the potential to study cognitive processing with high BOLD CNR to enable the detection of more subtle cognitive responses and/or the precise characterisation of responses on an individual subject level. However, it is expected that brain function in higher cortical regions may be more intrinsically variable between subjects than in primary sensory cortex due to the combination of inter-individual anatomical differences in brain structure and/or spatial differences in functional responses. Thus a method by which brain structure and function can be studied on an individual subject basis and linked to behaviour would be highly beneficial to the advancement of cognitive neuroscience.

Here, in order to investigate whole brain higher-order cognitive function at UHF (7T), an adapted Posner paradigm, a classic paradigm in cognitive neuroscience, is used. Typically, a Posner paradigm is used to modulate visual spatial attention (e.g. (Posner 1980, Gitelman, Nobre et al. 1999, Corbetta, Kincade et al. 2000, Gould, Rushworth et al. 2011)), and has been less commonly used to modulate spatial attention in the somatosensory domain (e.g. (Haegens, Handel et al. 2011, Haegens, Luther et al. 2012, Wu, Li et al. 2014)). In electroencephalography (EEG), these attention modulations have been associated with increased hemispheric lateralisation of the power of alpha-frequency (8-13Hz) oscillations over sensory-specific areas with increased spatial attention to a location (i.e. a contralateral decrease and ipsilateral increase in alpha power relative to the attention location) (Worden, Foxe et al. 2000, Rihs, Michel et al. 2007, Gould, Rushworth et al. 2011, Haegens, Handel et al. 2011, Haegens, Luther et al. 2012, Zumer, Scheeringa et al. 2014). This alpha modulation has been associated with a decrease in inhibition/increase in cortical excitability in the relevant cortical areas when attention is directed to its corresponding spatial location (Worden, Foxe et al. 2000, Rihs, Michel et al. 2007, Gould, Rushworth et al. 2011, Haegens, Handel et al. 2011, Haegens, Luther et al. 2012, Zumer, Scheeringa et al. 2014). In fMRI studies, Posner paradigms have been widely used to identify brain regions involved with visual spatial attention (Gitelman, Nobre et al. 1999, Martinez, Anllo-Vento et al. 1999, Corbetta, Kincade et al. 2000, Carrasco 2011), identifying modulations across a number of cortical regions including the frontal eye fields (FEF), posterior parietal, cingulate, striate and extrastriate cortex (Gitelman, Nobre et al. 1999, Martinez, Anllo-Vento et al. 1999), with the intraparietal sulcus (IPS) recruited specifically during the cued attention period prior to stimulus presentation (Corbetta, Kincade et al. 2000). Whilst tactile spatial attention has been studied less commonly with EEG and fMRI, the inferior parietal lobule and secondary somatosensory cortex (SII) have been shown to be recruited (Wu, Li et al. 2014, Gomez-Ramirez, Hysaj et al. 2016). To our knowledge, only one previous fMRI study has investigated the brain activity underlying manipulation of spatial attention across two sensory modalities in a single task using a Posner-style paradigm (Macaluso, Eimer et al. 2003). The authors showed that directing attention spatially to the right or left side of the visual domain or tactile (somatosensory) domain generated two forms of attentional brain response, termed “unimodal” and “multimodal”. Unimodal effects were found in regions where responses were only seen for attention to that specific modality; with the superior occipital and fusiform gyrus recruited by vision and the post-central gyrus by touch.

Multimodal effects were independent of the attended sensory modality, and even observed when no stimulus was presented (only attention directed); these effects were strongest in superior premotor areas and the left inferior parietal lobule, but also seen in posterior parietal and prefrontal cortices. These previous studies illustrate that these paradigms recruit top-down attentional modulation and involve higher cortical frontal-parietal areas as well as primary sensory regions, thus providing an ideal paradigm for testing and developing the utility of UHF fMRI for cognitive studies.

Here, we use a visual/somatosensory top-down attention modulation task to assess brain regions involved in varying the degree of attention directed to the somatosensory and visual domains. To our knowledge, the areas involved in both directing and modulating attention between sensory modalities, such that attention is divided between the modalities, are currently unknown and provide an excellent test of UHF fMRI in the identification of the higher-order cognitive areas, where only subtle differences in the amplitude of the BOLD response between conditions are expected. We hypothesize that: 1) linear modulations of BOLD signal by attention should be observed in primary sensory regions (akin to previously reported EEG modulations for directing spatial attention (Gould, Rushworth et al. 2011)); 2) sensory modality independent modulations of BOLD signal (i.e. only dependent on how attention is split between modalities) will be seen in higher-order cortical regions, such as parietal cortex and FEF (Macaluso, Eimer et al. 2003). The benefits and challenges of performing a large FOV, whole brain study of higher-order cognition at 7T are presented. We aim to demonstrate the optimal analysis methods to study whole brain focal, higher-order cortical responses at the group level, where differences in BOLD response between conditions are subtle and individual anatomical and functional variability are evident. fMRI data are analysed on the cortical surface at both the individual subject and group level. At the group level, normalization using both volume and surface registration is assessed, along with the dependency on spatial smoothing. We highlight the limitations of standard analysis pipelines for the assessment of cognitive paradigms using high spatial resolution fMRI data at UHF, and the applicability of optimised analysis methods for such UHF fMRI studies.

## Methods

### Subjects

This study was conducted with approval from the local ethics committee and complied with the Code of Ethics of the World Medical Association (Declaration of Helsinki). All subjects gave written informed consent. Data were acquired from 10 experienced fMRI subjects (age 28 ± 5 yrs (mean ± s.d.), 4 female).

### MRI acquisition procedures

All MR data were acquired on a 7T Philips Achieva MR scanner (Philips Medical Systems, Best, Netherlands), with head-only transmit coil and 32-channel receive coil (Nova Medical, Wilmington, USA). Foam padding was used to minimize head movement.

#### Attention Modulation scan session

To ensure whole brain coverage with high temporal and spatial resolution a multiband (MB) [or simultaneous multi-slice (SMS)] (GyroTools Ltd, Zurich, Switzerland) gradient echo echo-planar imaging (GE-EPI) sequence was employed (TR=1.9 s, TE=25 ms, 1.5 mm isotropic resolution, 128 × 131 matrix, multiband factor 2, 75°flip angle (FA), SENSE factor 2.5, receiver bandwidth 1172 Hz/pix, phase encoding (PE) direction: anterior-posterior). 58 contiguous axial slices covering visual, somatosensory and attention-related regions (parietal cortex, dorsal-lateral prefrontal cortex (DLPFC)) were collected in the given TR period. B_0_-field maps were acquired (TR=26 ms, TE=5.92 ms, ΔTE=1 ms, 4 mm isotropic resolution, 64 × 64 matrix, 40 slices, FA =25°, SENSE factor 2) and local image-based (IB) shimming performed, thus limiting field perturbations in B_0_ over the whole brain FOV in the fMRI acquisitions.

A total of 210 fMRI volumes were acquired per run, with 30s/80s of baseline data collected at the start/end of each run whilst subjects fixated on a dot. MB data were reconstructed offline (CRecon, GyroTools Ltd). During all fMRI scans, cardiac and respiratory traces were recorded for physiological correction. A peripheral pulse unit (PPU) on the subject’s left ring finger was used to record the cardiac trace, and a pneumatic belt placed around the chest was used to record respiration.

In the same scan session, a high-resolution whole brain phase-sensitive inversion recovery (PSIR) sequence (Mougin, Abdel-Fahim et al. 2016) [0.7 mm isotropic resolution, 288 × 257 matrix, 98 slices, TI=785/2685 ms, SENSE factors: 2.2 (right-left, phase encode), 2 (foot-head, slice selection)] was also acquired for segmentation and cortical flattening.

#### Retinotopic Mapping scan session

We also perform a retinotopic mapping task on all subjects to provide functional boundaries in primary visual cortex, allowing comparison of functional and anatomically defined boundaries in a primary sensory region. Retinotopy GE-EPI fMRI data were acquired in a separate scan session (TR=2 s, TE=25 ms, 1.5 mm^3^ isotropic resolution, 124 × 121 matrix, 85°FA, SENSE factor 2.5, receiver bandwidth 1089 Hz/pix, PE direction: foot-head). 32 coronal oblique slices were acquired to cover the entire visual stream (V1 to IPS), with IB shimming performed over this target region, and 120 volumes collected per run.

##### Paradigm

###### Attention Modulation paradigm

Subjects viewed a projector screen through prism glasses whilst lying supine in the scanner bore. A variant of the Posner paradigm (Posner 1980) was used to modulate attention between visual [V] and somatosensory [S] domains, as shown in Figure 1A. This comprised of a 250 ms duration visual cue at the start of each trial to indicate the certainty (0%, 40%, 60% or 100%) of the target appearing in the visual domain (Fig. 1A, cues panel). This was followed by a blank screen with a central white fixation dot, which was presented for an attention inter-stimulus-interval (aISI) of variable length of 1.3 - 1.6 s prior to stimulus presentation. During this aISI subjects were instructed to allocate their attention between the visual and somatosensory domains, according to the cue certainty. A target stimulus (high or low frequency) was then presented in either the visual or somatosensory domain, with a distractor stimulus (middle frequency) presented concurrently in the other sensory domain. The visual stimuli comprised a Gabor grating presented for 66.7 ms with spatial frequency of 3.2 (low), 6.4 (middle) or 12.8 (high) degrees/cycle which filled a visual angle of 2.1°in the lower left visual field (at 5.2/2.6 degrees of visual angle in the horizontal/vertical planes). The somatosensory stimuli consisted of piezoelectric stimulation at 4, 16 or 52 Hz presented for 250 ms to the tip of left index finger (Dancer Design, St. Helens, United Kingdom, http://www.dancerdesign.co.uk). The different stimulus durations between the visual and somatosensory stimuli were required to ensure comparable task behaviour responses across sensory domains (validated prior to the fMRI study). Subjects were required to respond as quickly as possible after stimulus presentation, with a button press of the right index or middle finger to indicate if the target stimulus was delivered at low or high frequency, providing accuracy scores and reaction time measures. An 850 ms period was allowed for subjects to respond to the stimulus presentation before the visual cue for the next trial was presented.

**Figure 1:**
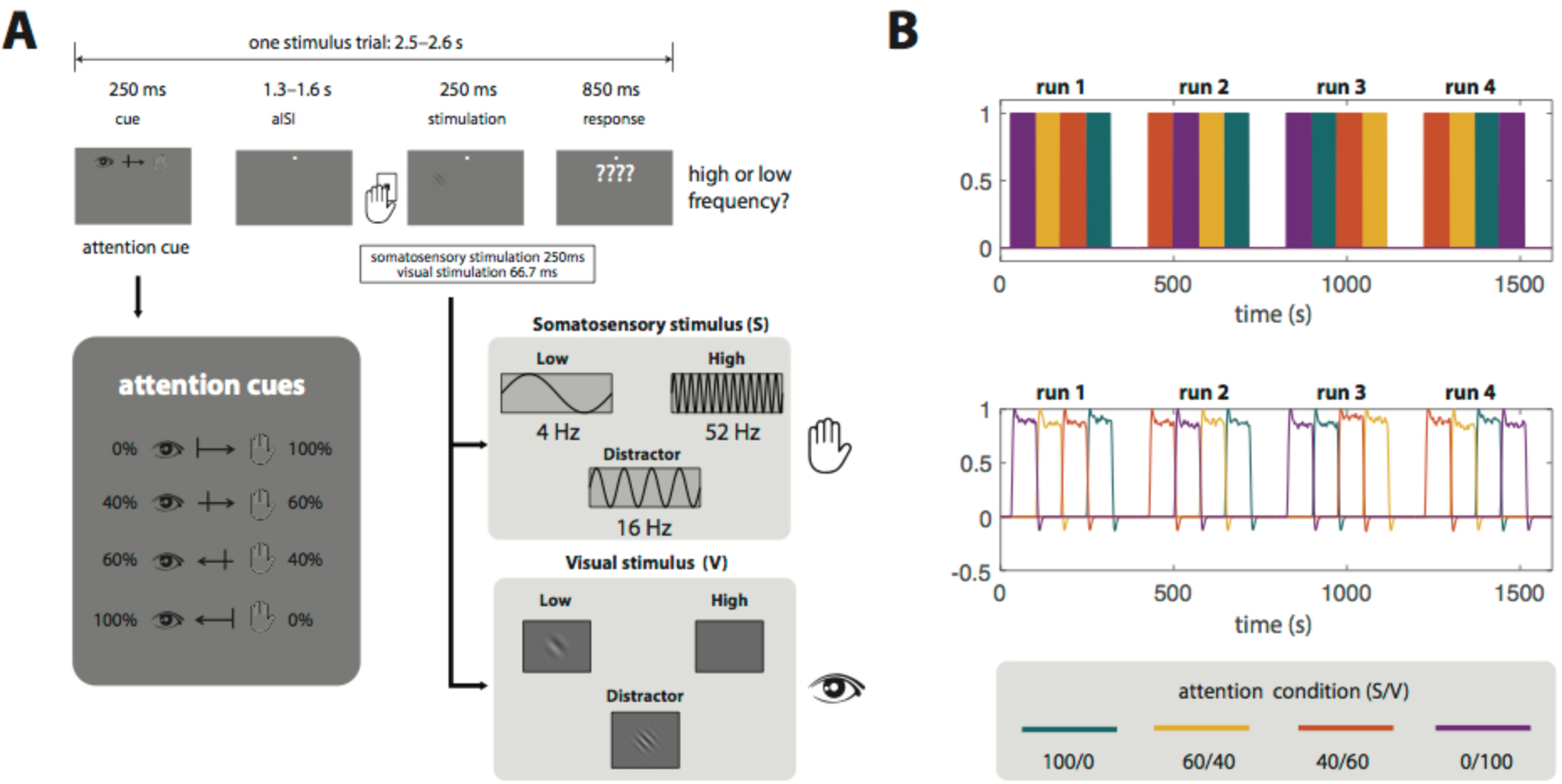
Schematic showing the attention paradigm. **A**: illustration of a single trial, inserts show all possible cue and stimulus presentations within a trial. **B**: Schematic showing full fMRI experiment. Trials (A) were presented in blocks of 25 of the same cue condition (e.g. 100% V) before switching to another cue condition (e.g. 40% V, as shown in run 1). Cues were presented in a pseudo-random order across runs. A 10 minute rest period (not shown) was provided between runs.

Trials were presented in blocks to ensure high sensitivity to the attention modulation. 25 trials of a given cue condition (S/V: 100/0, 60/40, 40/60, 0/100) were presented in a block before switching to a different cue condition. Within one fMRI run, blocks of each of the four cue conditions were presented in a pseudo random order. A total of four fMRI runs were acquired in a single scan session giving a total of 100 trials per condition over all runs (see schematic in Fig. 1B). Approximately 10 min rest between runs with no fMRI acquisition was provided (to provide a break for subjects and allow for scanner gradient cooling). At the end of each fMRI run the subject was given visual feedback to inform them of their performance (accuracy of target classification). The stimulus paradigm was controlled by Psych-toolbox (ptb v3 – http://psychtoolbox.org/overview/).

Prior to entering the MR scanner, subjects performed one run of the fMRI paradigm (i.e. 25 trials of each condition) to minimise learning effects during the fMRI acquisition and ensure they could perform the task well.

###### Retinotopic mapping paradigm

Eccentricity and polar angle maps were measured using standard retinotopic mapping procedures comprising an expanding annulus and rotating wedge to define visual areas (V1, V2, V3, V4) for each subject, akin to (Gardner, Merriam et al. 2008). These are standard retintopic stimuli provided in the mgl toolbox (version 2.0 https://github.com/justingardner/mgl <u>using mglRetinotopy.m</u>). Eccentricity was measured using an expanding annulus that started from a fixation point at the fovea and moved out to the periphery to map visual eccentricity. To measure polar angle in visual cortex, a wedge rotated clockwise. Both the annulus and wedge stimuli were textured with a checkerboard with alternating chromatic contrast. One period of stimulation (i.e. a full expansion from fovea to the periphery or a complete clock-wise rotation of the wedge) took 24 s, with ten repeats collected per scan. For both annuli and wedges, a second scan was collected with reverse order (i.e. from expansion to contraction, or clock-wise to counter-clockwise) to control for the spatiotemporal haemodynamic response function (Aquino, Schira et al. 2012). For all conditions subjects fixated on a central cross which flickered between red and grey.

## Analysis

### Behaviour

To test the efficacy of the paradigm in modulating attention, three-way repeated measures ANOVAs were performed on the accuracy and reaction time measures acquired during the fMRI task. Data were tested for significant effects of cue (i.e. 0/100% compared with 40/60% attention), modality (i.e. attending to the somatosensory or visual modality) and subject (i.e. whether behaviour was the same for all subjects). When significant interactions were found, post-hoc analyses using paired t-tests were performed to identify the conditions driving the observed differences.

### MRI Pre-processing

#### Functional MRI data

fMRI data were first corrected for physiological noise to remove cardiac and respiratory associated noise using retrospective image correction (RETROICOR) (Glover, Li et al. 2000). Images were motion corrected (SPM12, http://www.fil.ion.ucl.ac.uk/spm/software/spm12/) to the first GE-EPI volume within each fMRI run, and then between fMRI runs by registering all data to the mean GE-EPI image of the second fMRI run. The functional images were then corrected for scanner drift by regressing out a linear drift between the initial and final rest periods of each run.

#### Anatomical MRI data

PSIR data were processed to derive a bias-field corrected PSIR image (Mougin, Abdel-Fahim et al. 2016) by polarity restoring (using the phase) the first inversion time (785 ms) and dividing by the sum of the modulus of the two inversion time images (785 and 2685 ms). The PSIR images were then automatically segmented into grey an1d white matter using Freesurfer V6.0 [freesurfer.net], taking care to manually correct any segmentation errors. The two interfaces between grey/white matter and grey matter/cerebral spinal fluid (CSF) generated the *white* and *pial* surfaces respectively.

The mean GE-EPI image across all runs of a paradigm (attention or retinotopy) was then used to co-register from fMRI data space to PSIR data space using a linear affine transform (mrTools, http://gru.stanford.edu/doku.php/mrtools/overview).

### Subject normalisation

Subjects’ fMRI data were normalized into a standard space for group analyses. Both volume and surface based registration techniques were performed for comparison. These steps are summarized in Figure 2, with the processing pipelines described in detail below.

**Figure 2:**
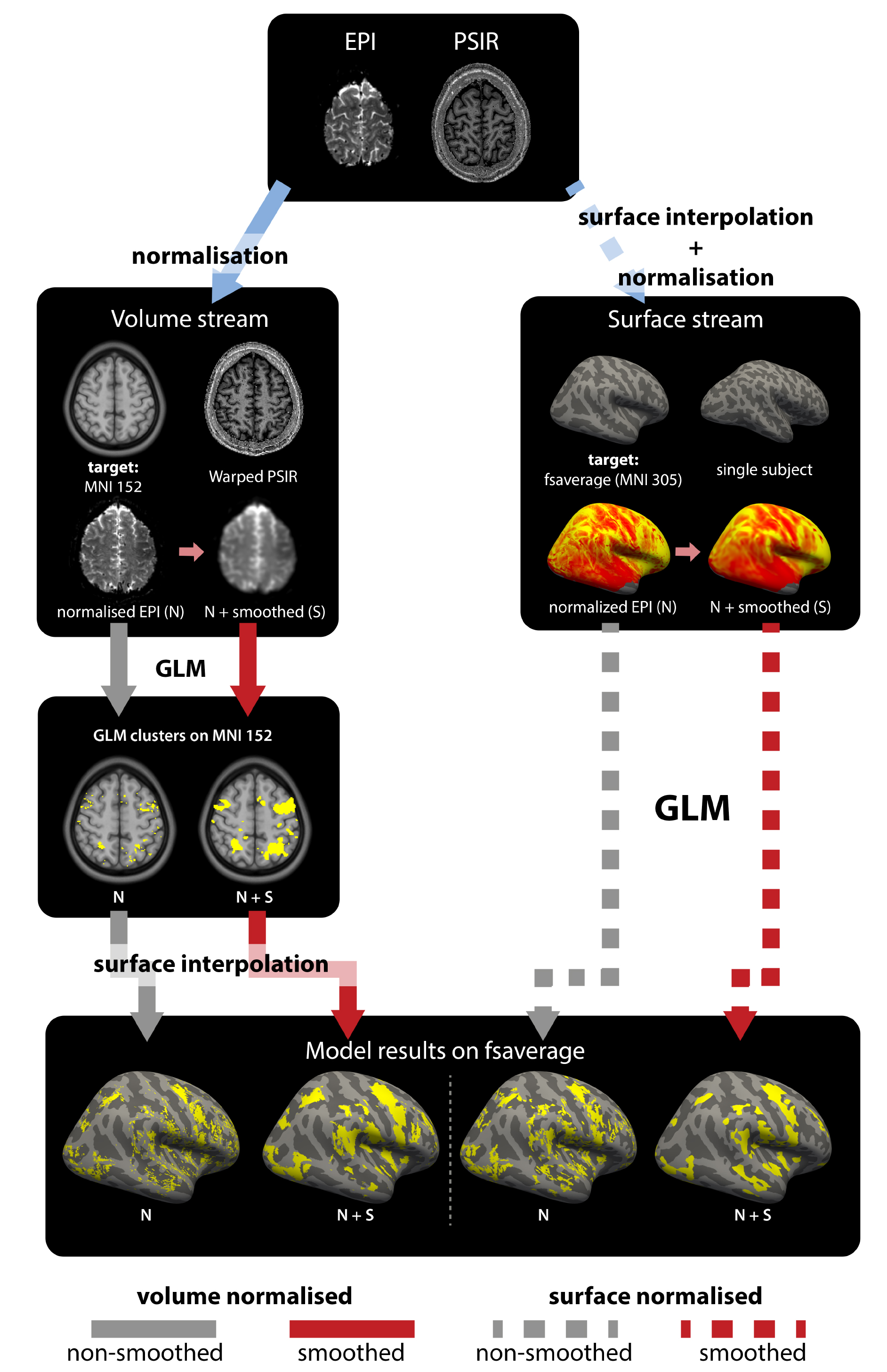
Schematic showing the main processing pipelines of fMRI data using volume normalisation (left stream) and surface normalisation (right stream). It should be noted that in both streams, there is a trilinear interpolation from volume to surface space. Clusters shown are those in response to the task for a single subject using a standard GLM, p<0.05 FWE. Note, that in the figure N denotes normalised EPI data and S denotes smoothed data.

#### Volume normalisation

For volume normalisation, subjects’ PSIR data were registered to the Montreal Neuroimaging Institute (MNI) 152 space using “unified segmentation” as implemented in SPM12 (http://www.fil.ion.ucl.ac.uk/spm/software/spm12/). Briefly, this normalization procedure was performed in a single generative model that involves affine co-registration to MNI 152 space and classification of tissue types using a Bayesian mixture model of tissue types. These warping parameters were then applied to the pre-processed fMRI data (see Fig. 2, solid lines path).

#### Surface normalisation

For surface normalisation, subjects’ PSIR data were registered to a surface generated from MNI 305, Freesurfer’s average subject template *fsaverage,* described in (Fischl, Sereno et al. 1999). In brief, registration was performed in two steps. First, the PSIR segmentations were inflated and the vertices from the individual subject surface were mapped onto an individual subject spherical representation of the brain and curvature information regarding the folding patterns of the gyri and sulci for the individual subjects derived (standard procedure in Freesurfer). The individual subject folding patterns were then used to register these surfaces to the normalized *fsaverage* surface, which had been mapped onto a sphere, as previously described (Fischl, Sereno et al. 1999). Second, fMRI data were interpolated onto the individual subject cortical surface and a “white” layer, pial layer, and mid-layer (at 50% of the surface normal between the white and pial layer) were defined using *mri_vol2surf* in Freesurfer. This resulting fMRI data was then normalised to *fsaverage* using the transform defined from the PSIR data (see Fig. 2, dashed lines path).

#### Visualisation of normalised data

Note, for visualisation of volume normalised data, the interpolation step described above for surface normalisation (moving the white, pial, and mid-layers onto the surface) was also performed. Data from both normalisation streams were additionally displayed on a flattened representation of fsaverage, calculated by making cuts to the cortical surface and using a cost function to metrically optimize (in terms of distance) the flattened representation (available as standard in Freesurfer). A tool to aid visualization of the flattened maps, used in this manuscript, is freely available at (https://github.com/KevinAquino/freesurferFlatVisualization.git).

### Smoothing of functional data

fMRI data in both volume and surface normalised space, were spatially smoothed (Fig. 2, smoothed (S) shown by red lines) using a kernel of FWHM of 4.5 mm. In the volume stream, this is equivalent to applying a 3D Gaussian kernel; in the surface stream, the data were smoothed across the surface over a ring that corresponds to a kernel of 4.5 mm diameter at each vertex.

### functional MRI post-processing

#### Individual subject General Linear Model (GLM) Design

Since the main focus of this work was to localise brain regions recruited by top-down modulation, the attention period (aISI) during each trial was modelled in the GLM for each subject. All trials were modelled regardless of behavioural response given the high performance to this task. For each of the four conditions (i.e. one combination of S/V), the aISI time periods of each trial were set up as blocks and convolved with the standard haemodynamic response function for each relevant software package (SPM or Freesurfer), Fig. 1B, bottom panel. This was repeated across the four fMRI runs resulting in 16 model estimates as regressors in a 1^st^ level design matrix. In addition, the motion parameters were included as covariates of no interest and a constant term for each run to model differences in baseline GE-EPI signal between runs.

#### Functional contrasts

Contrasts were assessed in a 1^st^ level analysis and individual subject beta-weight (*β*) values computed. The contrasts assessed are summarised in Figure 3. They comprised: (i) a task contrast (Fig. 3B) weighting all attentional conditions equally – to probe any brain regions recruited by the task independent of attentional cue; (ii) a positive linear contrast (Fig. 3C) – to identify regions whose BOLD response linearly co-varied with increasing visual attention; (iii) a negative linear contrast (Fig. 3D) – to identify regions whose BOLD response linearly co-varied with increasing somatosensory attention; and (iv) a task difficulty contrast (Fig. 3E), using an “n” shape to weight the “hard” conditions (S/V 40/60 or 60/40) more than the “easy” conditions (S/V 100/0 or 0/100) – to identify regions whose BOLD response was modulated by the division of attention between modalities, independent of where the attention was directed.

**Figure 3:**
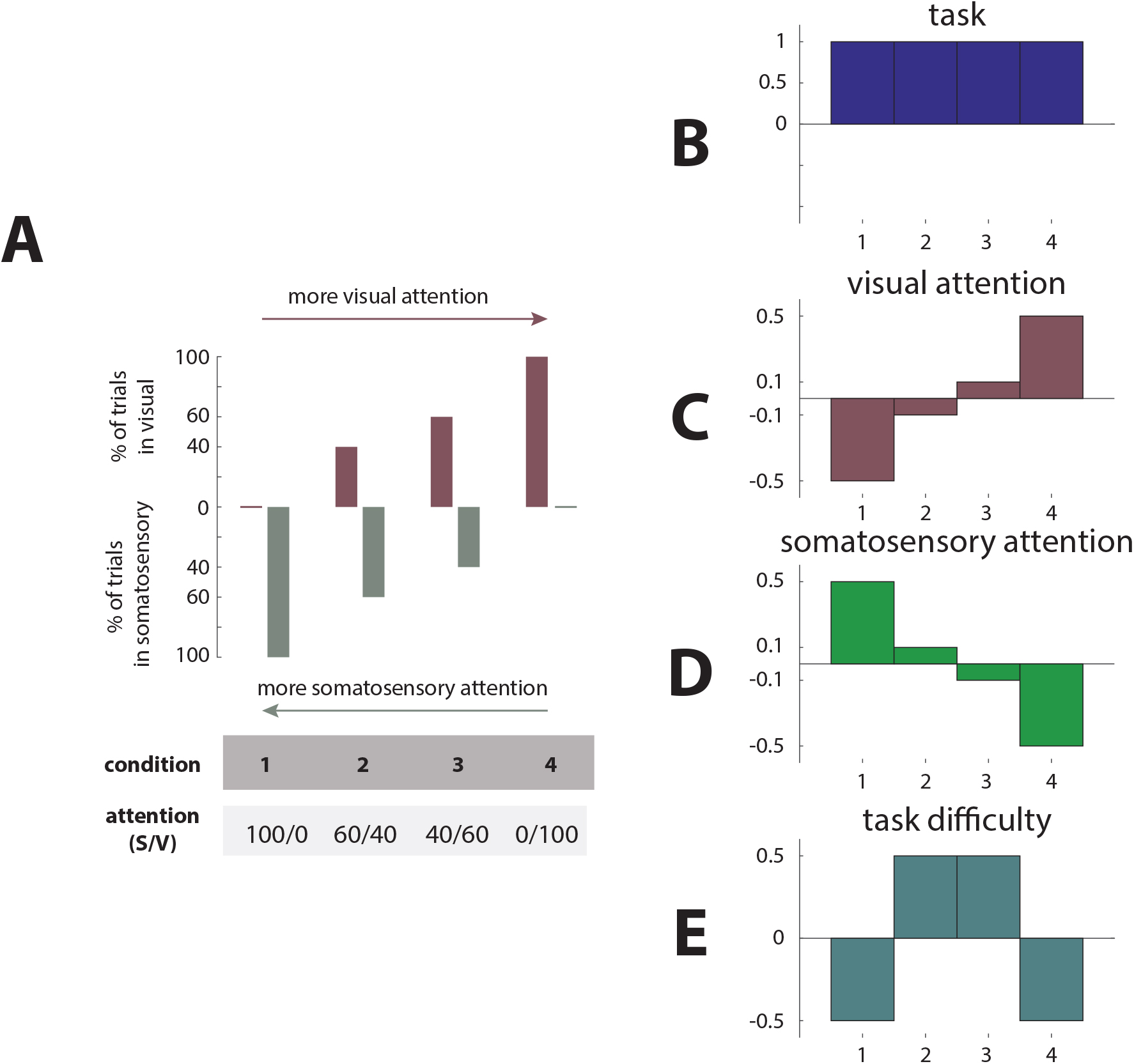
Schematic showing the different contrast conditions used in the 1^st^ Level GLM analysis **A.** The % of trials in the somatosensory/visual domain for each of the conditions. **B**-**E** show the different contrasts used in the GLM analysis to interrogate the effects of attention.

A one-sample t-test was used to threshold the maps for each functional contrast, which were then corrected for multiple comparisons. For volume normalised data, this correction was calculated using random field theory via the formulation of resolution elements (RESEL), as implemented in SPM12 (Worsley, Marrett et al. 1996). For surface normalised data, RESELS are not simply described analytically (Hagler, Saygin et al. 2006), thus Monte Carlo simulations were used to generate an equivalent estimation (Hagler, Saygin et al. 2006), as implemented in Freesurfer v.6.0.

#### Atlas Definition

To interrogate functionally specific regions, two functional cortical atlases were used. The Glasser atlas (as shown in Figure 4) comprising 180 regions per hemisphere, based on data from multimodal imaging (HCP-MMP 1.0)(Glasser, Coalson et al. 2016), and the Freesurfer Destrieux Atlas (2009) (Destrieux, Fischl et al. 2010). These two atlases were applied to the normalized *fsaverage* brain (Freesurfer v 6.0) (see “Subject normalisation”) and on individual subject surface reconstructions for spatial interrogation of fMRI responses. Figure 4 shows the flattened representation, with region labels, that is used throughout the results and discussion sections.

**Figure 4:**
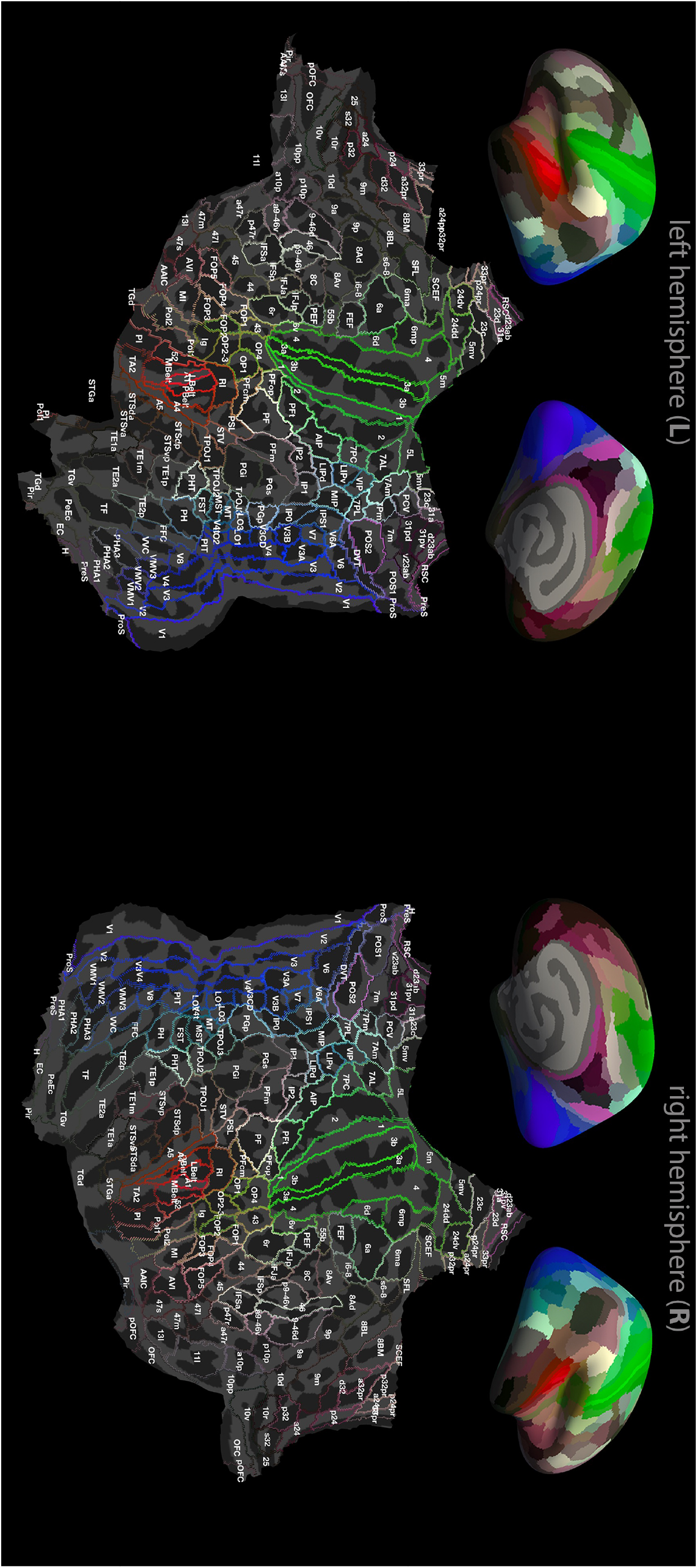
The relationship between the whole inflated brain (top row) and the Glasser atlas overlaid on the flattened patches (bottom row) – note colours on the inflated brains correspond with the line colours on the flattened patches demarcating anatomical boundaries. Labels written on the flattened patches are taken from the Glasser atlas (Glasser et al. 2016).

### Group level fMRI analyses

Functional results were analysed at the group level using three different methods. Methods 1 and 2 were designed to interrogate the data allowing for the expected spatial variability in activation due to differences in anatomy and/or function between subjects. Method 3 performed a standard 2^nd^ level GLM group analysis.

Method 1 was performed in both volume and surface normalised space, for both unsmoothed and smoothed data (see Fig. 2). This method combined the corrected (p<0.05, FWE corrected) one-sample t-test maps from each individual subject to form functional inter-subject conjunction maps. First, surviving voxels/vertices in the 1^st^ level maps were used to form binary maps for each subject. These binary maps were then summed resulting in functional inter-subject conjunction maps ranging from 0 to 10 (representing each subject).

Method 2 involved analysis of the surface normalised, unsmoothed data using the 180 parcellated regions defined by the Glasser atlas (Glasser, Coalson et al. 2016). Within each parcel and for each subject, the vertices with the top 5% of t-statistical values in response to the ‘all’ task condition were found to create the region of interest for that subject. The average *β* values for each condition for the vertices within the region of interest computed. These *β* values were then normalized across conditions (using the maximum *β* value of any condition) for each subject. Performing this analysis in surface normalised space ensured that the same number of data points were used per subject for a given region. The dependence of the normalized *β* values on visual attention (or decreasing somatosensory attention) was then modelled for each of the 180 parcellated regions. Three candidate models were tested to match contrasts used in the GLM analyses (as shown in Fig. 3): a constant model (task), a linear model (modality specific attention) and a negative quadratic (task difficulty) model. The ‘winning’ model was selected to be that which minimized the Bayesian Information Criterion (BIC) – a metric that “rewards” model fit and “punishes” model complexity (Schwarz 1978). The resulting analysis in the Glasser atlas regions within a given hemisphere were then corrected for false discovery rate (FDR) at a q-factor 0.1.

Method 3 performed a standard 2^nd^ level group GLM analysis on the volume normalised smoothed data (see Fig. 2, red solid lines). The summary statistic from each contrast for the 1^st^ level GLM analysis was used to perform a one-sample t-test in a conventional 2^nd^ level mixed effects GLM analysis.

### Assessing whole brain fMRI data quality

Temporal signal-to-noise (tSNR) was used as a measure of quality of the functional imaging data by calculating the mean of each voxel’s fMRI time series divided by its standard deviation. The voxel-wise calculation of tSNR, was performed after the fMRI time series were corrected for linear and quadratic drift (motion was < 0.5 mm in all subjects). tSNR measures were computed both before and after RETORICOR to determine the spatial improvements afforded by physiological correction. tSNR maps were interpolated onto the cortical representation for each subject on the surface template *fsaverage*.

### Retinotopy analysis

The data from the two retinotopic paradigms: the rotating wedges and the expanding/contracting annuli, were used to map visual polar angle and eccentricity, respectively. The data were processed with standard retinotopic analyses using mrTools (http://gru.stanford.edu/doku.php/mrtools/overview). Following motion correction and co-registration, the scans from the wedge paradigm were combined: first, scans from both the clockwise and counter-clockwise condition were shifted by 2 frames, then the order of the volumes of the scans from the counter-clockwise condition were reversed prior to averaging with the scans form the clockwise condition. This reversal and shift was used to cancel out the effects from the spatiotemporal haemodynamic response function (Aquino, Schira et al. 2012). Following this average, the time series at each point were correlated with a cosine function with frequency that matched the stimulus delivery. The analysis provides a correlation – which indicates model fit, and a phase angle – which was correlated to the phase when the stimulus was presented and thus visual polar angle. Voxels that survived a correlation threshold of 0.4 were analysed for their phase. Half the visual hemifield contained phases that ranged from [0,π] whereas the other half ranged from [π,2π]. Boundaries where the phase reversed were interpreted as borders of visual areas (Engel, Rumelhart et al. 1994, Schira, Tyler et al. 2009). A similar procedure was repeated for the annuli paradigm, where the phase maps [0,2π] were used as additional validation of a visual area. The retinotopic maps were visualized on surface and flattened representations (as detailed in the “Normalisation” section) and used as a validation of the functional relationship to the structural, atlas based, cortical parcellation used.

## Results

### Subject Behavioural Performance

Figure 5 shows the group mean behavioural responses to the task and indicates clear modulation of task performance, both accuracy and reaction time, across cue conditions. We show a significant (p<0.05, three-way repeated measures ANOVA) effect of the cue condition on both accuracy (p=9×10^−9^; F=400.1) and reaction time (p=4.9×10^−10^; F=771.5). Lower accuracy (Fig. 5A) and longer reaction times (Fig. 5B) were observed for the trials when subjects divided their attention (40/60 and 60/40 cue conditions) compared to when subjects focused their attention on one modality (0/100 and 100/0 cue conditions). We also observed a significant effect of modality on accuracy (p=1.9×10^−4^; F=36.5) and reaction time (p=5×10^−9^; F=456.1), such that somatosensory attention modulated reaction times by a greater amount than visual attention. This was reflected by a significant cue × modality interaction for reaction time (p=1.4×10^−8^; F=362.2), with significantly shorter reaction times to somatosensory stimuli than visual stimuli (Fig. 5B). In addition, a significant difference between subjects (p=0.014; F=4.8 and p=5.5×10^−4^; F=4.8, repeated measures ANOVA) in behavioural measures (accuracy and reaction time) was observed, demonstrating that all subjects did not perform the task equally, with some finding the paradigm more difficult than others.

**Figure 5:**
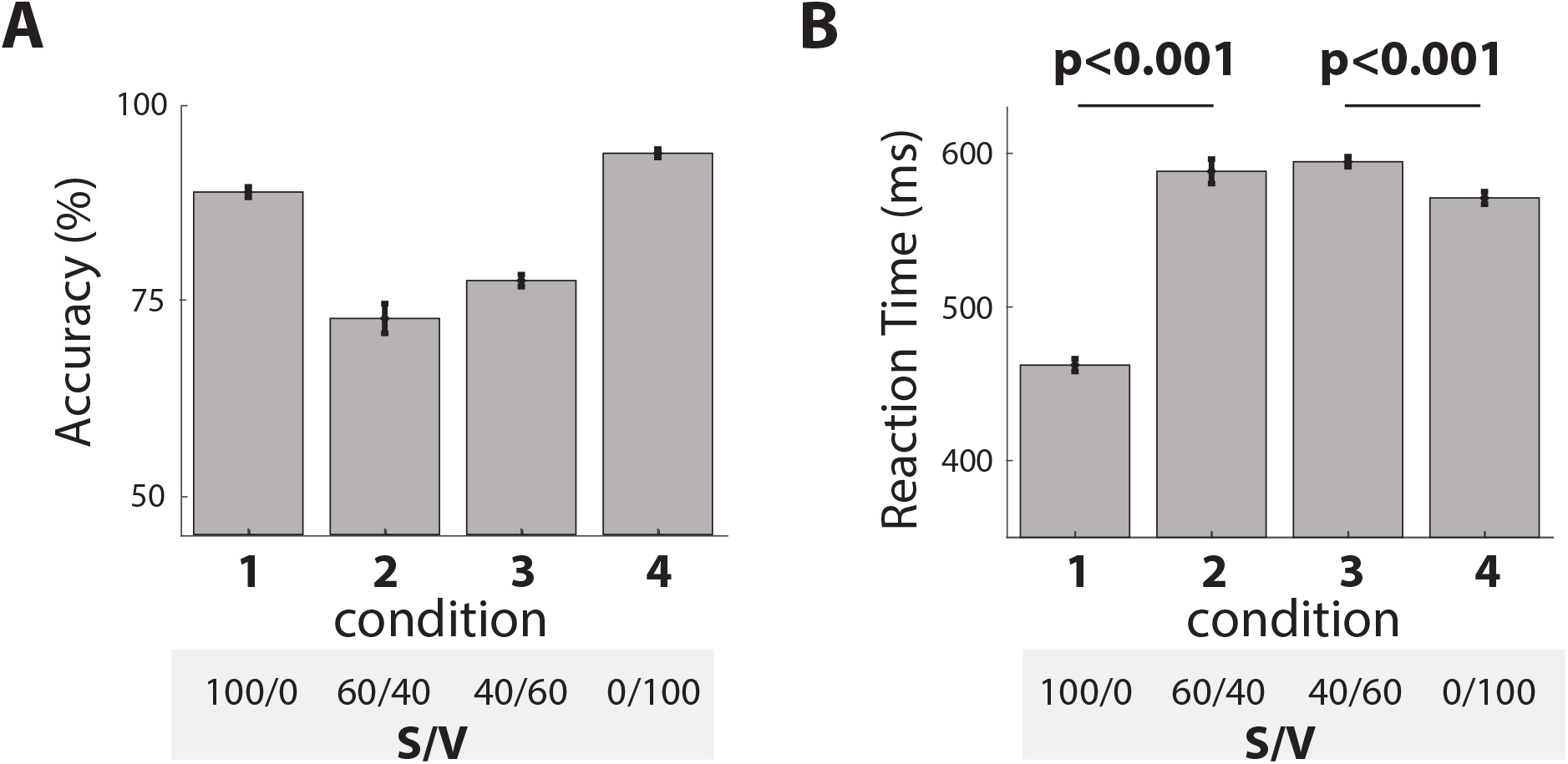
Group mean behavioural responses for (**A**) accuracy and (**B**) reaction time. A significant (three-way repeated measures ANOVA, p<0.05) effect of cue, modality and subject on both accuracy and reaction time was observed. An interaction of cue and modality was observed for the reaction time, post-hoc t-test analyses reveal the differences (marked on **B**). Error bars indicate SEM.

### Assessing whole brain fMRI data quality

tSNR was assessed for the attention fMRI data over the whole cortex and shown to be relatively homogeneous (Fig. 6). Data showed the characteristic lower tSNR in the temporal lobes due to higher physiological noise (Hutton, Josephs et al. 2011), and in this region physiological correction provided the greatest benefits (Fig. 6B). Lower tSNR was also observed within the central sulcus, this was driven by a reduced mean signal, likely caused by the heavy myelination in this region (Glasser and Van Essen 2011). Focal thin lines showing a reduction in tSNR adjacent to the pre- and post-central gyri are likely to be an artefact of the cortical unfolding process.

**Figure 6:**
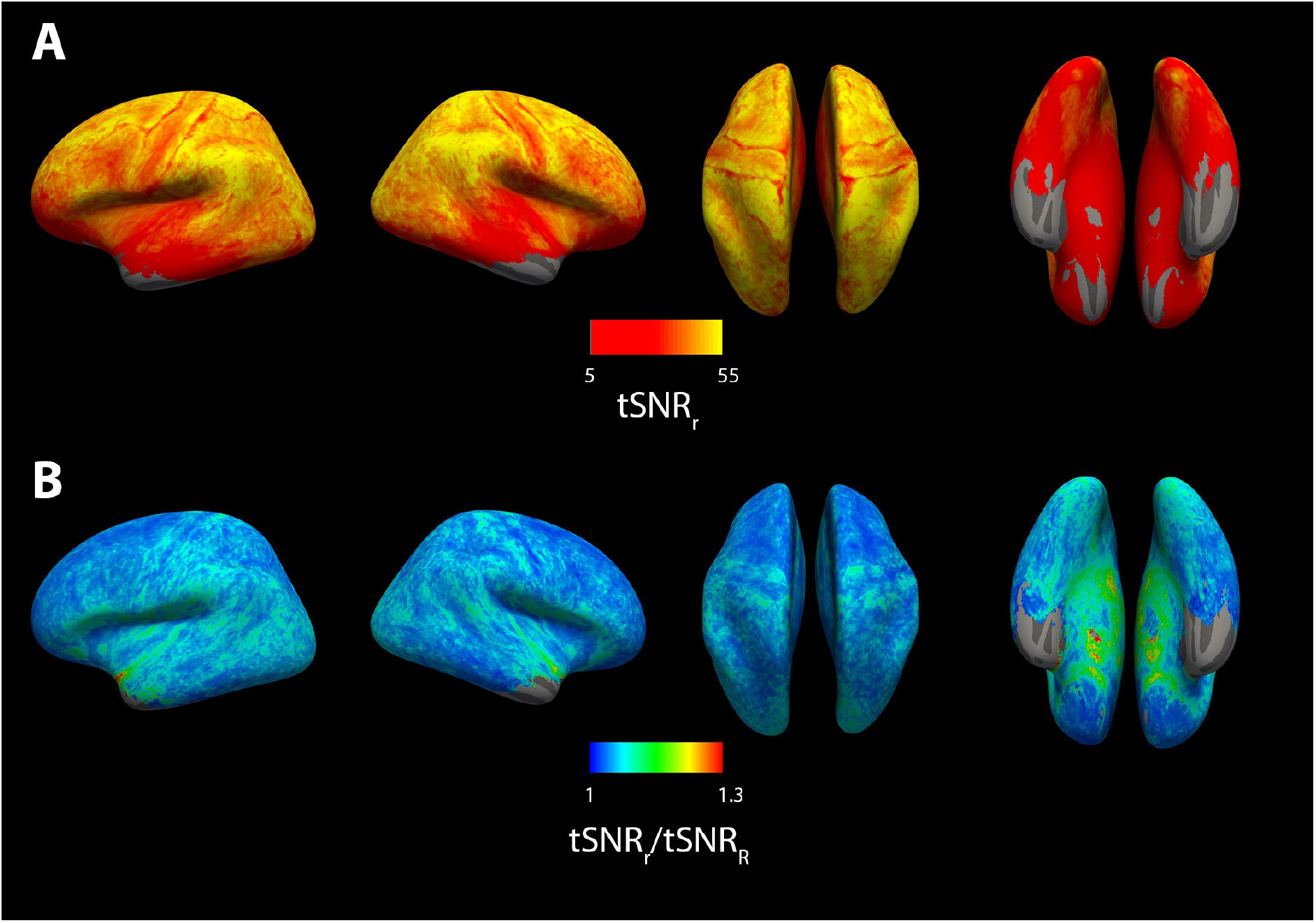
**A**: tSNR maps for the attention task fMRI data after physiological noise correction. **B**: Maps of the ratio of tSNR_after correction_/tSNR_before correction_ showing those areas with the greatest increase in tSNR following physiological noise correction.

### Anatomical variance

Figure 7 illustrates the greater inter-subject anatomical variability in both size and location of higher-order cognitive areas such as the IPS (Fig. 7A), compared with the primary sensory regions such as the primary visual cortex (Fig. 7B). Figure 7A and B show the atlas definitions of IPS and V1 on individual surfaces respectively, and the surface registered folding patterns (as indicated by the signed mean curvature K – a proxy for Sulci and Gyri as shown on the colorbar of Fig. 7A&B). Across subjects, for the atlas defined IPS region, differences can be seen in the pattern of gyri (blue) and sulci (red) included with the region (highlighted with arrows in Fig. 7A, bottom row). In particular differences of the folding patterns inside, and in the neighbourhood of, the automated definition. The high quality of the surface normalisation procedure for all subjects in the primary visual cortex (V1), where there is less anatomical variability, can be verified by our retinotopy data (Fig. 7C). This provides a functional map of visual region boundaries which show strong spatial agreement with the Glasser atlas defined anatomical ROIs.

**Figure 7:**
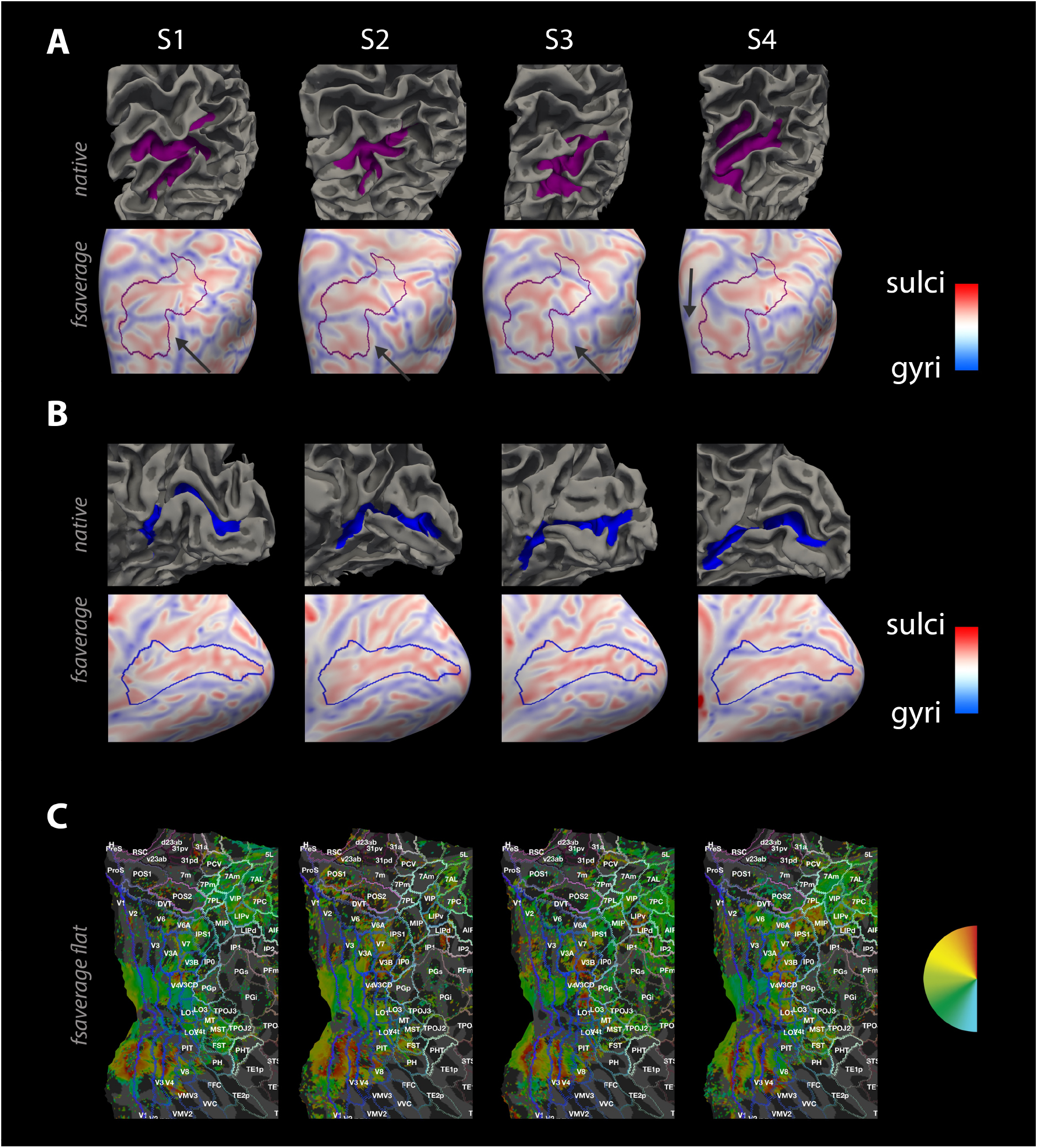
**A:** Right IPS on native individual subject (S1-S4) surfaces (upper row, purple region shows IPS delineated by Freesurfer’s registration method and Freesurfer Destrieux atlas), and on normalized space (lower row, IPS outlined in purple) where the individual subject curvature has been mapped into this space. Note the variability in the spatial pattern of sulci and gyri across subjects: there are clear differences in the anatomy (sulci shown in red, gyri shown in blue) within the IPS boundary (purple line) between subjects, as highlighted by the arrows. **B:** Data as for **A** shown for primary visual cortex (V1) demarcated in blue. Note the greater agreement of the sulci (shown in red) within V1 area across subjects in the normalized images. **C** Individual retinotopy on normalized flattened surface with the Glasser atlas overlaid. Note the retinotopy phase-reversals (corresponding to the left visual hemifield shown in the semicircle) denoting functional boundaries map onto the anatomically defined visual regions (V1-V3).

### Individual response to attention task

At the individual level, all subjects showed a significant (p<0.05, FWE corrected) response to the task contrast condition (Fig. 3B) and modulation to the cue condition, with the largest and most extensive responses generally seen for the task difficulty contrast condition (Fig. 3E) with greater activation observed for the 40/60 conditions than the 0/100 conditions. However, the extent and location of the functional activation for each of the contrasts varied spatially across subjects relative to the structural information defined by the Glasser atlas (Fig. 4), even when surface normalisation was used. Figure 8 illustrates this in two representative subjects, showing that whilst activation clusters to a given contrast were observed in approximately the same region, there was not exact spatial agreement across subjects, likely due to anatomical differences as well as the degree of functional response to the task.

**Figure 8:**
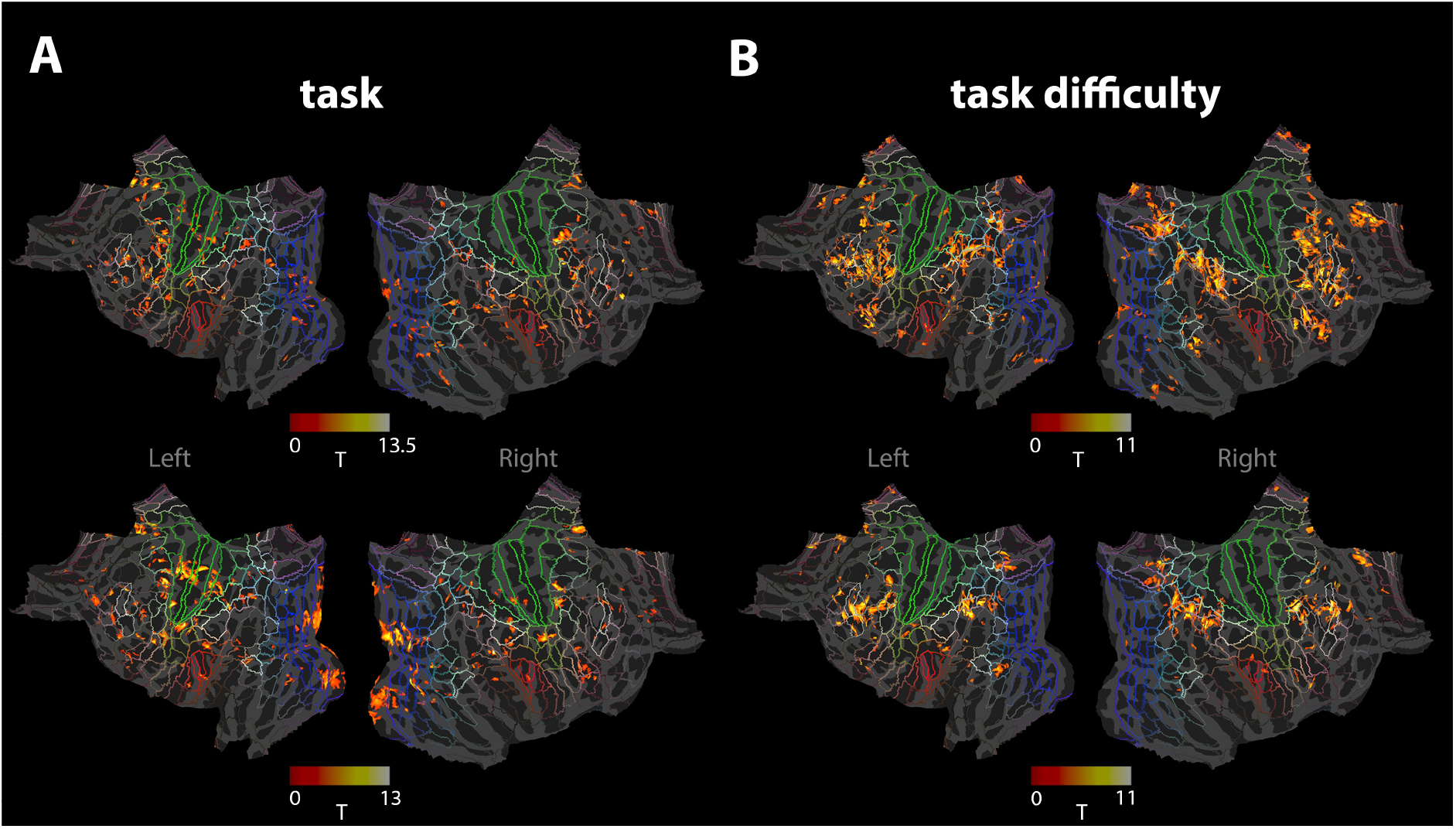
Maps showing regions of significant (p<0.05, FWE corrected) activation to the *task* contrast (**A)** and *task difficulty (***B**) contrast (see Fig. 3B and 3E) for two subjects (top and bottom row, respectively). Individual subject maps have been surface normalised to the fsaverage template. Lines demarking anatomical regions are derived from the Glasser atlas, as shown in Figure 4.

### Surface versus volume normalisation

The effect of the spatial variability in functional responses is highlighted when considering the normalisation of responses to a standard template and the effect of smoothing the data. Figure 9 compares the effect of volume and surface normalisation on the spatial agreement of significant areas of activation in response to the attention task paradigm across subjects. The importance of the choice of normalisation method can be seen by comparing the spatial overlap conjunction map of frontal-eye-field (FEF) and intraparietal sulcus 1 (IPS1) activations for volume (Fig. 9B &D) and surface (Fig. 9F &H) normalisation (see Fig. 4 for region definitions). Volume normalisation results in poor inter-subject spatial agreement, with the overlapping activity in 3 or less subjects in FEF (Fig. 9B) and IPS1 (Fig. 9D), due to the focal nature of the responses combined with the observed anatomical variability in higher order cognitive areas (Fig. 7A). A much greater spatial overlap was observed for surface normalisation (Fig. 9A, F and H), with overlapping activity in up to 7 subjects in the FEF (Fig. 9F) and 5 in the IPS1 (Fig. 9H).

**Figure 9:**
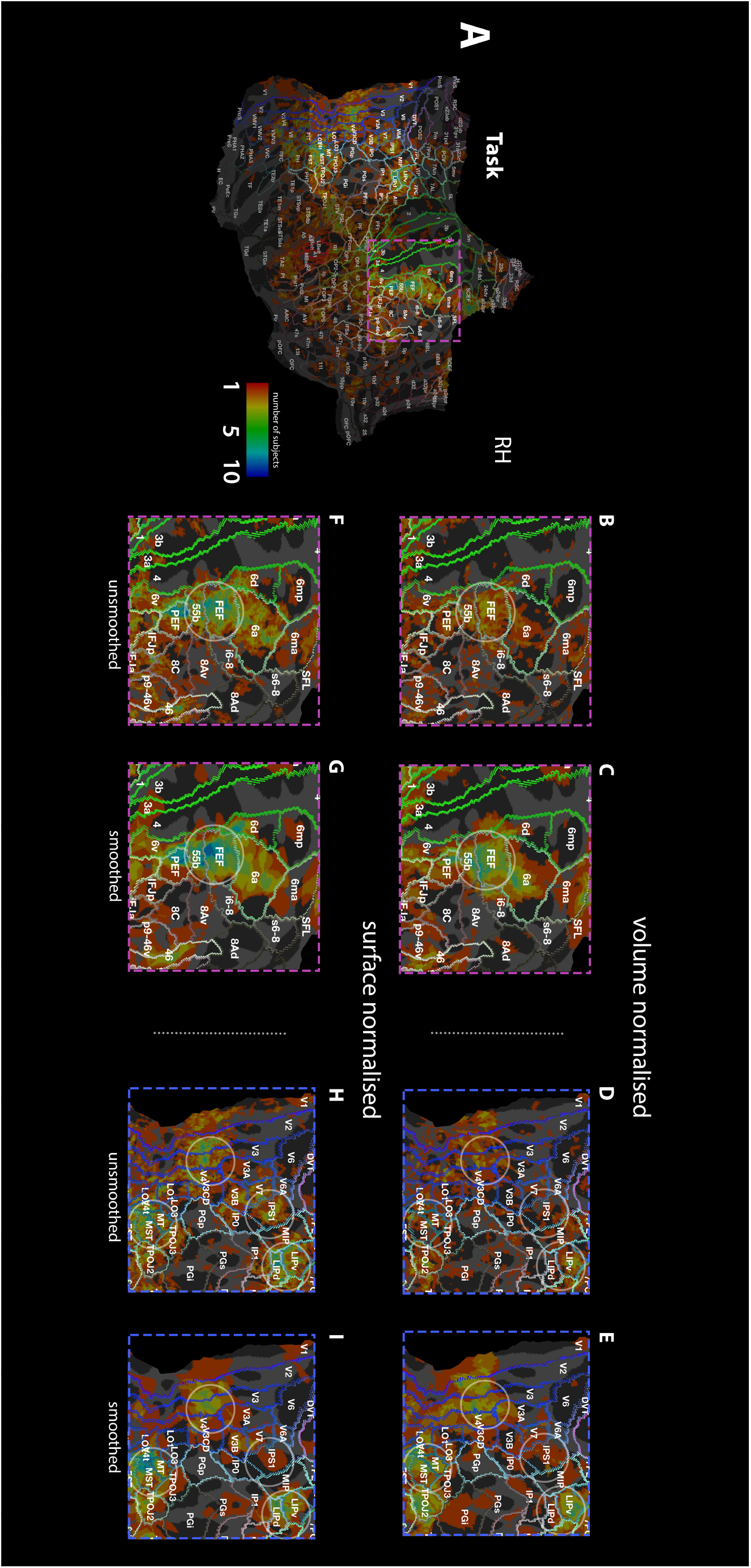
Comparison of group conjunction of individual subject T-stat maps (p<0.05, FWE corrected) for the *task* contrast between different pre-processing pipelines. **A**: group conjunction map for the whole right hemisphere, created from the surface normalisation and no smoothing stream (see Fig. 2, grey dashed line). Pink box includes frontal-eye-field area shown in Panels **B&C+F&G** whilst blue box includes the intraparietal cortex shown in Panels **D&E+H&I**. Panels **B&D** show volume normalisation without spatial smoothing. Panels **C&E** show the volume normalisation with 4.5 mm FWHM spatial smoothing. Panels **F&H** show surface normalisation without smoothing, whilst Panels **G&I** show surface normalisation with 4.5 mm FWHM surface smoothing. Circles in Panels **B-I** draw attention to regions where conjunction of T-stat maps varies greatly dependent on the processing pipeline employed. For similar maps for task difficulty condition and visual attention condition see Figure S2.

Spatial smoothing also has profound effects on the inter-subject overlap of activations, as can be seen by comparing Figure 9B/D with Figure 9C/E for volume normalisation, and Figure 9F/H with Figure 9G/I for surface normalisation. Applying spatial smoothing to the volume normalised data (Fig. 9C&E) helped compensate for anatomical variability, especially in the FEF, but the resultant spatial agreement was still lower than for surface normalisation alone. Spatial smoothing of the surface normalised data reduced the spatial overlap in some areas such as the IPS1 and V3 (Fig. 9G&I), since focal responses will be reduced when a smoothing kernel of greater extent than the activity is used. Similar effects were observed for the *task difficulty* contrast. Figure 10 illustrates the advantage of surface normalisation (Fig. 10B) over volume normalisation (Fig. 10A) for the detection of both the *task* contrast and *task difficulty* contrast across a group (see Fig. S1 for inferior views of Fig. 10). Figures 9 and 10 show that in general, response to task was observed in the visual stream from V1 to V4 and into the lateral intraparietal (LIP) region as well as in the FEF, whilst task difficulty correlated with activity of the IPS and dorsal lateral prefrontal cortex (also see Fig. S2 top). In addition, activations to the positive linear contrast (attention to visual domain) were observed, but these were far more focal than the task condition or task difficulty condition, resulting in little inter-subject spatial overlap (Fig. S2 bottom), even when using the optimal pipeline of surface normalisation and no spatial smoothing. No significant activations to the negative linear contrast (attention to somatosensory domain) were seen consistently over the group, regardless of analysis stream.

**Figure 10:**
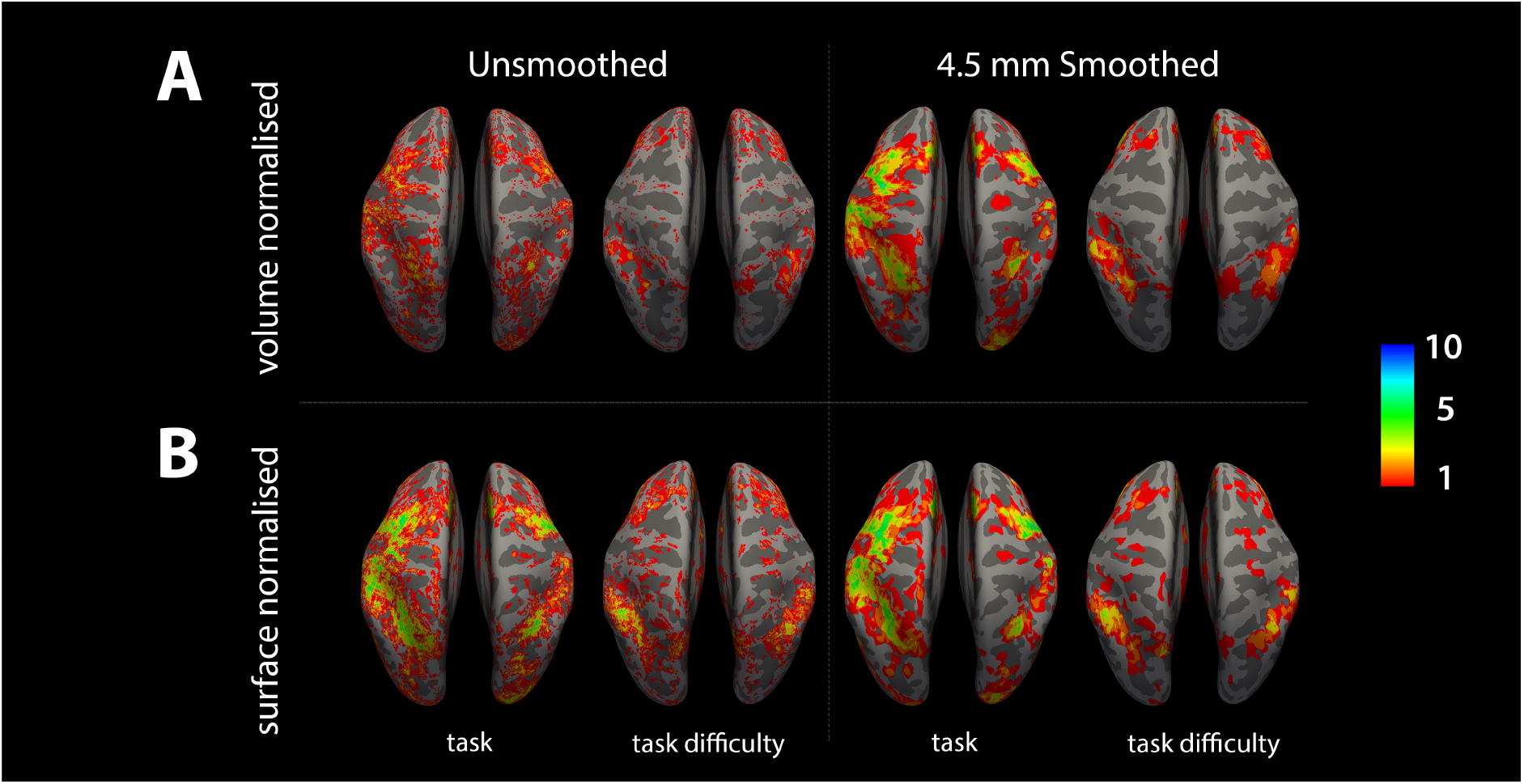
Group conjunction of individual subject T-stat maps (FWE p<0.05) showing number of subjects with overlapping activation using the different processing pipelines **A:** volume normalisation, **B:** surface normalisation. For A) and B) results are shown for unsmoothed (left) and smoothed (4.5 mm FWHM kernel) data (right). For Inferior views, see Figure S1.

### Assessment of attentional contrasts

Figure 11 shows the result of performing model fits (constant, linear or quadratic) to the individual subject normalized *β* values, across all four conditions, for each anatomical region of the Glasser atlas. This provided an alternative method to investigate those areas which showed a significant linear (i.e. increased visual or somatosensory domain attention) or “n”-shape modulation (i.e. task difficulty) in response to the attention task. For a given anatomically defined region, statistical tests were performed only on voxels that responded to the task contrast (Fig. 3B) in that parcel for each individual subject. A significant “n”-shape correlation was found in a number of regions (Fig. 11A&B1, blue), with large clusters of parcellated regions seen in the parietal and dorsal lateral prefrontal cortex. Significant linear correlations all showed a positive gradient (i.e. increased visual attention) and were seen in the visual cortex and VIP (Fig. 11A&B, red-orange). The most robust linear modulations were seen in the Fusiform face complex (FFC) (p = 0.008, FDR corrected) with the strongest linear modulations of the lower-order visual areas (V1-V4) found in V4 (p = 0.06, FDR corrected) [V3: p = 0.09, FDR corrected]. The strongest “n”-shaped modulation was seen in area 7m (p = 0.002, FDR corrected) of parietal cortex and in Area 44 (p = 0.005, FDR corrected) of dorsal lateral prefrontal cortex. An example region of interest, V3, and its model fit are shown in Figs 11C&D, respectively.

**Figure 11:**
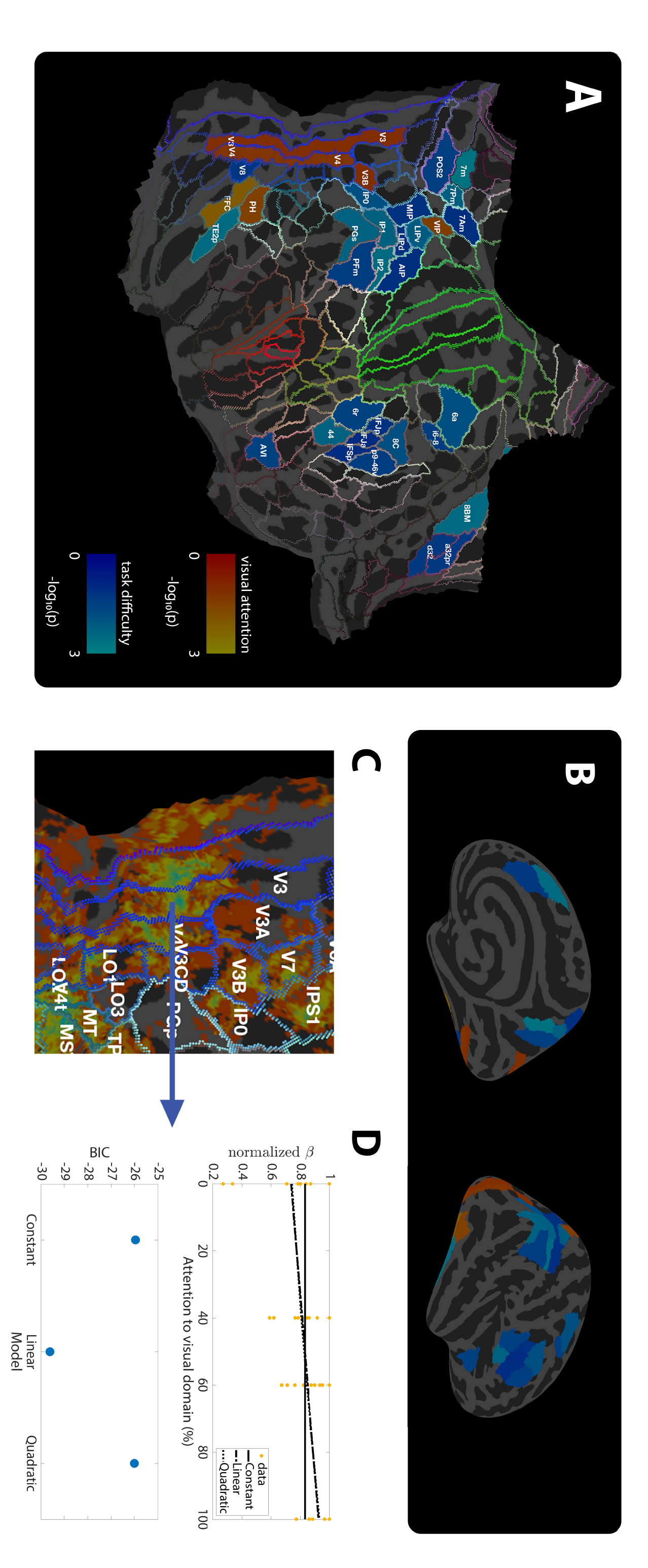
**A:** The winning models (linear or quadratic, assessed with BIC) calculated across cortical regions (FDR adjusted and threshold at p<0.1); highlighting the regions showing linear modulation (red/orange- i.e. increased visual or somatosensory domain attention) and quadratic modulation (blue - i.e. “n”-shape task difficulty). **B:** Inflated views of the data shown in A. **C:** selected region of interest (V3) with group conjunction of response to task (akin to Fig. 10A). **D:** An example of a model fit (top panel) to the top 5% of b values from region V3, and the BIC calculated for the three candidate models, showing a linear fit clearly “winning” in this region.

### Standard group analysis

Figure 12 shows standard mixed effects group analysis results for the attention task, highlighting that standard methods do not readily reveal the modulations of the functional responses to this cognitive task even when using a relatively lenient threshold of p<0.001, uncorrected. Focusing on the right hemisphere responses, as stimuli were presented to the left; we observed that, whilst some areas showed the response to task regardless of the attentional cue (Fig. 11, dark blue), only very small disparate areas showed activation to the task difficulty (quadratic, shown in pale blue) or attention to a modality (linear, positive – shown in red, negative – shown in green) across the conditions. This lack of response was due to lack of spatial agreement across subjects (see Figs. 7, 9 and 10) and highlights the need for optimal analysis pipelines to investigate such responses in high spatial resolution fMRI data. Additional activations observed in the left hemisphere are likely to be related to the button press response and the response to the task, rather than the direction of attention and therefore were not the focus of this study.

## Discussion

We explored the feasibility of using high spatial resolution UHF fMRI to interrogate the BOLD response to a cognitive task across the whole brain. First, we investigated the magnetic field inhomogeneity (∆B_0_) and the temporal SNR of the data, showing high quality whole brain data was achieved. We then explored the inter-subject differences in brain structure in both primary visual areas and higher-order cortical regions and its likely contribution to inter-subject differences in the spatial location of functional responses. We showed that considerable differences in anatomy are present in higher order cortical areas, whilst primary visual regions showed good anatomical agreement, as published previously (Fischl, Rajendran et al. 2008).

Given the observed inter-subject structural and functional differences, we investigated the effect of normalisation and smoothing procedures on the spatial agreement of the functional response to the cognitive task, in particular in higher order areas. We show that the choice of normalisation and smoothing procedures employed is critical when the fMRI response of interest is focal and/or in higher-order cortical regions. We demonstrate a novel group fMRI analysis for assessing attentional modulation to the cognitive task, by fitting *β* weights derived from 1^st^ level GLM analysis to different models within each parcellated region. We reveal areas which show quadratic and linear modulations of BOLD signal in response to the attention conditions of the task, findings which are not clearly observed using standard 2^nd^ level mixed effects GLM analysis due to the large inter-subject spatial variability and difference in amplitude of functional response modulations.

### Data quality, anatomical variability and parcellation of brain regions

When considering whole brain functional responses, it is important to first consider the GE-EPI data quality. We used IB-shimming and B_0_ mapping to provide in good global B_0_ homogeneity for the attention fMRI data acquisition, this ensured a subvoxel shift of pixels in the GE-EPI data in the phase encode direction compared to the anatomical data. In future we will improve this further with the use of dynamic distortion correction techniques (Visser, Poser et al. 2012). tSNR was also relatively homogeneous over the cortex, though there was a noticeable reduction in the temporal lobes, regions known to be most greatly affected by physiological noise (Hutton, Josephs et al. 2011), and the central gyrus, driven by a reduced mean signal caused by heavy myelination in this region (Glasser and Van Essen 2011).

Group analysis of fMRI data typically requires data normalisation to interrogate all subjects’ data in the “same space” and to allow the study of responses in equivalent regions. Given the known anatomical variability between subjects (Geyer, Weiss et al. 2011), it is questionable whether normalisation of data to a standard template is appropriate using any transformation (i.e. volume or surface normalisation). However, a method of parcellating the brain, or defining anatomical structures, is necessary to form group comparisons. In the absence of individual subject sub-millimetre anatomical scans of multiple MR contrasts for parcellation of brain structures (Tardif, Schafer et al. 2015), surface normalisation with a detailed anatomical atlas, such as the Glasser atlas (Glasser, Coalson et al. 2016), may be the best approach.

Surface normalisation approaches match the curvature of the sulci and gyri of individual subjects to a template. In a recent study, Tardif *et al* demonstrate that surface based methods can be refined to improve spatial normalisation based on such curvature (Tardif, Schafer et al. 2015). However, they also highlight that it may not be beneficial to maximise normalisation based on curvature and cortical folds, since in higher cortical areas, the curvature may not reflect the functional boundaries of regions (Tardif, Schafer et al. 2015). Instead, they suggest information is required to allow alignment of individual brains based on functional boundaries. These functional boundaries are commonly believed to be reflected by cytoarchitecture, but it is not possible to interrogate cytoarchitecture directly *in vivo.* Instead, T_1_ maps may provide additional information on myelination, which is believed to closely relate to cytoarchitecture, to inform group normalisation (Turner and Geyer 2014, Tardif, Schafer et al. 2015) or allow parcellation of individual brain regions (Geyer, Weiss et al. 2011, Turner and Geyer 2014). However, the success of this form of normalisation has primarily been assessed on myelin rich primary sensory cortices. Whilst some gains have been highlighted in higher order regions, FEF and ventral intra-parietal area [VIP] (Tardif, Schafer et al. 2015), it remains to be assessed as to how useful this normalisation is for other higher cortical regions, where myelination is generally lower with less contrast between regions. Indeed, it is unclear whether whole brain normalisation based on T_1_ maps may bias warping of the brain to correctly align regions of high myelin at the expense of functional areas with low myelin. Our data highlights the importance of future developments to map functional boundaries in individual subjects *in vivo,* to enable subject-specific structural and functional parcellation (Geyer, Weiss et al. 2011, Robinson, Garcia et al. 2017). Such boundary alignment may also take account of boundaries identified from robust fMRI tasks, such as individual subject resting state and visuotopic maps, as employed by Glasser *et al* (Glasser, Coalson et al. 2016). With such subject specific parcellations, *β* weights could be extracted from individual subject anatomical/functional regions and fed into a fitting process as used here (Turner and Geyer 2014). Alternatively, the landmarks of individual subject borders may be used to provide a more accurate normalisation, as suggested by Tardiff *et al* (Tardif, Schafer et al. 2015), subsequently allowing more standard GLM approaches to be employed. However, it should be noted that such additional measures to define structural or functional boundaries in an individual subject come at the expense of considerable addition scan time.

### Spatial smoothing

Spatial smoothing of fMRI data is widely adopted at lower field strength to blur inter-subject structural differences in brain anatomy for group analyses, increase statistical power (Turner and Geyer 2014), and ensure data meets Gaussian Random Field theory assumptions for statistical analysis (Worsley and Friston 1995). Until recently, many UHF whole brain studies have employed considerable spatial smoothing (Boyacioglu, Schulz et al. 2014, Goodman, Wang et al. 2017, Mestres-Misse, Trampel et al. 2017).

However, recent papers (e.g. (Stelzer, Lohmann et al. 2014, Turner and Geyer 2014, Turner 2016)) highlight that the use of large smoothing kernels negates the benefits of the high spatial resolution of fMRI achievable at UHF. In addition, they highlight that smoothing is not required for False Discovery Rate correction (Turner and Geyer 2014) due to the inherent smoothness of fMRI data due to the point-spread function of the BOLD response (Stelzer, Lohmann et al. 2014) (Polimeni, Renvall et al. 2017). Turner (Turner and Geyer 2014) provides a detailed critique of the problems associated with spatial smoothing (Stelzer, Lohmann et al. 2014, Turner and Geyer 2014, Turner 2016). Here, we show the limitations of spatially smoothing high resolution fMRI data, highlighting that focal responses in higher-order areas can be diluted by smoothing to the point that these responses no longer survive statistical analyses, resulting in reduced overlap of activations across subjects (compare Fig. 9H and Fig. 9I).

Without spatial smoothing, methods to best deal with differences in brain anatomy become vital to minimise inter-subject spatial variability, especially for higher-order cognitive areas for which anatomical variability is greater than for primary sensory areas (Turner and Geyer 2014). Minimal anatomical variability in primary sensory cortex may, in part, explain the successes of UHF high spatial resolution studies of sensorimotor and visual cortex (Sanchez-Panchuelo, Besle et al. 2012, Goncalves, Ban et al. 2015, Sanchez Panchuelo, Schluppeck et al. 2015, Sanchez Panchuelo, Ackerley et al. 2016, Kemper, De Martino et al. 2017, Poltoratski, Ling et al. 2017, Rua, Costagli et al. 2017). Indeed, our data confirmed this observation, showing good inter-subject correspondence of V1 (Fig 7B) and excellent correspondence of the anatomical and functional boundaries of V1-V3 (Fig 7C). However, anatomical agreement of higher-order areas, such as the IPS, was much poorer, with considerable variability across subjects observed even with the use of surface normalisation (Fig 7A).

### Limitations of group GLM analyses for cognitive tasks

Stelzer *et al* (Stelzer, Lohmann et al. 2014) have previously highlighted conceptually that fundamental differences in spatio-temporal representations of brain function leads to potential pitfalls when using a mixed effects GLM group analysis. They highlight that if the spatio-temporal pattern of response does not overlap completely across subjects, only a subset of the true activation for each individual will be present in the group analysis, i.e. the region where there is spatial agreement over subjects. This is due to both inter-subject anatomical differences and differences in functional brain activity due to the subjects’ response to the task (i.e. relationship to the canonically modelled response) which is likely to be particularly prevalent in cognitive tasks where individual subject strategy may differ. As such, a standard 2^nd^ level mixed effects GLM can only ever provide a partial picture of the true functional response to a cognitive task, as has also previously been shown from ICA and MVPA analysis (Etzel, Zacks et al. 2013, Xu, Potenza et al. 2013).

Our results (Fig. 12) corroborate the concerns raised in Stelzer’s thought experiment, demonstrating that a standard 2^nd^ level GLM analysis results in little common activation observed to any contrast (response to task, attention to visual stimuli, attention to somatosensory stimuli or task difficulty). This is due to the lack of precise spatial agreement between subjects (Figs 9, 10 & S1), despite use of a liberal p<0.001 uncorrected threshold and spatial smoothing. Whilst the extent of responses are increased using a fixed effects analysis (Fig S3), with a response to both the task contrast and task difficulty contrast observed (dark blue and pale blue), no linear modulations are seen. Therefore we propose that standard GLM approaches are not best suited to studies where functional responses are likely to vary across subjects due to task complexity and different task completion strategies.

**Figure 12:**
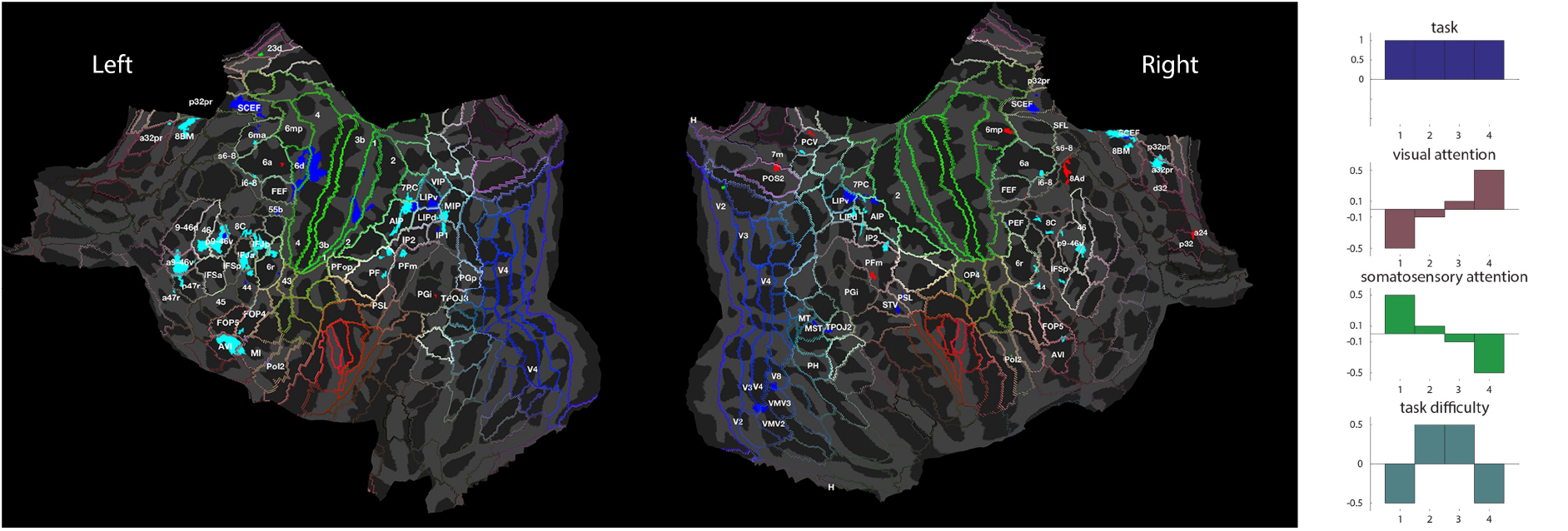
2^nd^ level GLM analyses of the volume normalised, smoothed attention data. Mixed effects analysis (p<0.001, uncorrected) activation maps (see Figure S3 for fixed effects analysis). Colours denote regions where the t-statistics for each of the contrasts exceeded the stated threshold, and were classified as “activated” regions to that contrast, with different contrast and associated colours shown in right hand panel.

### Functional interpretation of responses to attention paradigm

Tasks focussed on the direction of spatial attention to somatosensory stimuli have previously elicited responses in the inferior parietal lobe (IPL), FEF and SII (Wu, Li et al. 2014). Similarly the FEF and IPS, as well as posterior parietal cortex, cingulate, striate and extrastriate cortex (Martinez, Anllo-Vento et al. 1999, Corbetta, Kincade et al. 2000) have been shown to be active in response to visual spatial attention (Martinez, Anllo-Vento et al. 1999, Corbetta, Kincade et al. 2000). The attention related activations in the parietal and DLPFC regions (Fig 12) to sensory modality that we report using our optimised analysis pipeline are in line with previous observations. Here, we advance previous studies by varying the relative direction of attention between the visual and somatosensory domain, creating four conditions, whereas previous fMRI work has only directed attention solely from one location to another (spatial or modality). Such BOLD signal modulations between graded levels of attention are more subtle, requiring the higher CNR afforded by UHF fMRI.

We show that in parietal and DLPFC regions (Fig 12) the modulation related to attention level is quadratic, such that the fMRI BOLD response during the attention period is larger when attention is split between the two modalities (40/60 conditions) than when the attention is directed purely to one modality (0/100 conditions). This suggests that this attention effect is independent of the modality to which attention is to be directed and is instead related to task difficulty. This concept agrees with Macalauso et al (Macaluso, Eimer et al. 2003), who report modality independent modulations in superior premotor areas, left inferior parietal lobule, posterior parietal and prefrontal cortices, and (Corbetta, Kincade et al. 2000) which attributes activity in the IPS to be purely related to the top-down process of attention, rather than the response to a stimulus.

Our paradigm also allows us to differentiate regions that are independent of the modality to which attention is directed, from those regions where the modulation of the BOLD response is dependent on the modality that attention is directed to. We observe a linear modulation, increasing BOLD fMRI signal with increasing visual attention, within extrastriate visual cortex areas of V3, V4, V3b, area Parietalis (temporo-occipital) Basalis (PH, (von Economo and Koskinas 1925, Triarhou 2007, Glasser and Van Essen 2011)), Fusiform face complex (FFC) and VIP. Previously, analogous linear modulations of brain activity with attention have been reported in EEG data where alpha power has been shown to linearly decrease in the occipital/parietal regions of the hemisphere to which increasing spatial visual attention has been paid (Gould, Rushworth et al. 2011). However, EEG does not have the spatial resolution to identify the precise anatomical region in which the alpha power modulation is observed.

Of the lower order visual areas (V1-V4), V4 showed the most robust linear modulation with attention. Invasive animal electrophysiology recordings provide compelling supporting evidence that our analyses are identifying neuronal modulation by attention, since in these studies spike-spike coherence in the alpha and gamma bands has been shown to be significantly modulated by directed spatial attention in V4, but not in V1 (Buffalo, Fries et al. 2011). These invasive recordings showed that alpha power decreased when attention was directed to the visual area from which neuronal responses are measured. This report of a reduction in alpha power (Buffalo, Fries et al. 2011), negatively correlates with the observed increase in V4 BOLD response with increasing visual attention that was observed in this study, a finding supported by many previous electrophysiology reports of anti-correlation between alpha power and BOLD signals (e.g. (Goldman, Stern et al. 2002, Laufs, Holt et al. 2006, Mayhew, Ostwald et al. 2013, Mullinger, Mayhew et al. 2014)). It should be noted that the invasive recordings (Buffalo, Fries et al. 2011) also showed a concordant increase in gamma power in V4 but no significant gamma power change in V1. Previous reports show that gamma oscillations are generally thought to be most closely coupled to the BOLD response (Logothetis, Pauls et al. 2001, Magri, Schridde et al. 2012), suggesting that gamma changes could be driving the observed BOLD modulations we report. To our knowledge there are only reports of linear modulation of alpha with the graded manipulation of attention during a pre-stimulus cue period (e.g. (Gould, Rushworth et al. 2011)) but equivalent studies of gamma responses have not yet been performed.

Interestingly, we observed no negative linear modulations of BOLD responses across conditions (reflecting increased attention to the somatosensory domain), contrary to what might have been expected in the secondary somatosensory system (Wu, Li et al. 2014). Modulations in reaction time for attending 100% compared with 60% were larger when subjects attended to the somatosensory domain, than when attention was directed to the visual domain. Furthermore, the modulation of the accuracy measure between 100% and 60% was similar for both domains (no significant cue × modality interaction). Therefore the behavioural results strongly suggest it is unlikely that the subjects’ attention to the somatosensory domain was not modulated by this task, despite the lack of BOLD response in the somatosensory brain area. Modulation of modality specific alpha power with spatial attention have previously been reported for both the visual and somatosensory system (e.g. (Gould, Rushworth et al. 2011, Haegens, Handel et al. 2011, Haegens, Luther et al. 2012, Zumer, Scheeringa et al. 2014)), suggesting the processes behind directing spatial attention are not different for the two systems (visual and somatosensory). Further investigation is required to clarify the lack of responses in the somatosensory system to this type of attention paradigm where attention is divided between two modalities, rather than spatially.

Future studies may also benefit from the use of multi-variate pattern analysis (MVPA), which, by using spatial pattern recognition, has the potential to overcome some of the limitations of group GLM analyses (Turner and Geyer 2014). The optimal strategies presented here, i.e. surface normalisation and no spatial smoothing, should be considered to be complementary, providing an initial method to identify regions of interest on which MVPA can be performed. Furthermore, the methods presented in this study allow analyses on a smaller data set without the requirement for training and subsequent test data sets which can be challenging to obtain for complex cognitive tasks.

## Conclusion

This study shows the potential of 7T to study whole brain individual subject BOLD fMRI responses to a cognitive task. The optimal strategy of surface normalisation, no spatial smoothing and the analysis of responses within defined parcellations is demonstrated to assess cognitive processing involved in directing attention between sensory domains.

## Acknowledgements

The authors thank Olivier Mougin for his advice on PSIR images, Denis Schluppeck on advice on IPS, Sally Eldeghaidy for assisting with the scanning, and P.A. Robinson for his support during KA’s time in Australia. The work was funded by the Leverhulme Trust [grant number RPG-2014-369].

